# Organoid culture media containing growth factors of defined cellular activity

**DOI:** 10.1101/422923

**Authors:** Manuela Urbischek, Helena Rannickmae, Thomas Foets, Katharina Ravn, Marko Hyvönen, Marc de la Roche

## Abstract

The media components necessary for deriving and sustaining organoids from a number of epithelial tissues such as prostate, colon, gastric, liver, pancreas, and others have been established (1). Critical components of organoid media formulations are a set of growth factors that include EGF, R-spondins and BMP signalling antagonists such as Noggin or Gremlin. The practical limitation to organoid culture and the development of new applications for the technology is the use of defined cellular activities of growth factors in media formulations, in particular Noggin/Gremlin 1 and R-spondin 1. Here we report the production of highly pure recombinant Gremlin 1 and R-spondin 1 from bacterial expression and their use for culturing organoids. We detail the workflow for their purification, determination of cellular activity, quality control and their formulation in organoid media. The protocols we provide for generation of precisely formulated, cost-effective, organoid media of defined cellular activity will enable broader access to organoid technology and engender the development of novel applications.

## Introduction

The development of organoid technology has proven to be scientifically transformative with tremendous potential for application to the fields of epithelial biology and pathology and for clinical applications such as the generation of patient-derived organoid biobanks (2, 3). The critical component of organoid technology is the formulation of culture media with essential growth factors, in particular - R-spondin family members that potentiate Wnt pathway activity and BMP pathway inhibitors such as Gremlin 1 or Noggin 1 that inhibit differentiation cues. Currently, the production of growth factors for organoid media relies on eukaryotic expression systems to ensure correct disulphide bond formation and macromolecular folding. Typically, in eukaryotic expression systems, growth factors are secreted into the cell culture media from the expression hosts, and this conditioned media, containing the growth factor of choice (and all other proteins those cells naturally secrete), is then diluted directly into organoid culture media. However, R-spondin 1 and Gremlin 1/Noggin 1 expressing cell lines are not widely available and the use of conditioned media presents the problems of batch-to-batch variation of growth factor activity. Moreover, the presence of proteins, including growth factors, from serum present in cell culture media as well as those secreted from those cells may impact organoid growth. Commercially available purified growth factors can also be used to supplement organoid media however, these are expensive for medium to large scale applications such as genetic and chemical library screening or clinical biobanking of organoids. Moreover, commercial growth factors preparations can retain impurities from conditioned media that prevent accurate and reproducible determination of their cellular activities.

Here, we report a protocol for the generation of highly pure recombinant Gremlin 1 and R-spondin 1 from bacterial expression that contain negligible levels of endotoxin and free from impurities endemic to production using eukaryotic expression systems. We detail protocols for the determination of the cellular activities of the growth factors enabling the generation of media of reproducible potencies for organoids culture circumventing the issue of batch-to-batch variation. The production of defined and reproducible organoid media potencies has broad application for the culturing of organoids derived from a number of tissue types.

## Materials and Methods

### Expression plasmids

The pR-spondin 1 construct for production of R-spondin 1 is based on a previously reported pETDuet-based vector (4) for dual expression of a R-spondin 1-containing fusion protein (maltose binding protein, a tetra-cysteine tag, a thrombin cleavage site, residues 21-145 of human R-spondin 1 (Uniprot: O60565) and a hexa-histidine tag) and the *E. coli* disulphide bond isomerase, *DsbC.* We have inserted the coding sequence of Avi-tag upstream of the R-spondin 1 transgene by digestion of the original construct with *BamH*I and *Not*I and Gibson assembly to create pR-spondin 1 (Fig. 2A).Plasmids for testing R-spondin 1 activity in the potentiation of Wnt pathway activity are the Super 8x TOPFlash (SuperTop; Addgene 12456) and the *Renilla* Luciferase control reporter vector pRL-SV40P (Promega).

**Figure 1.**
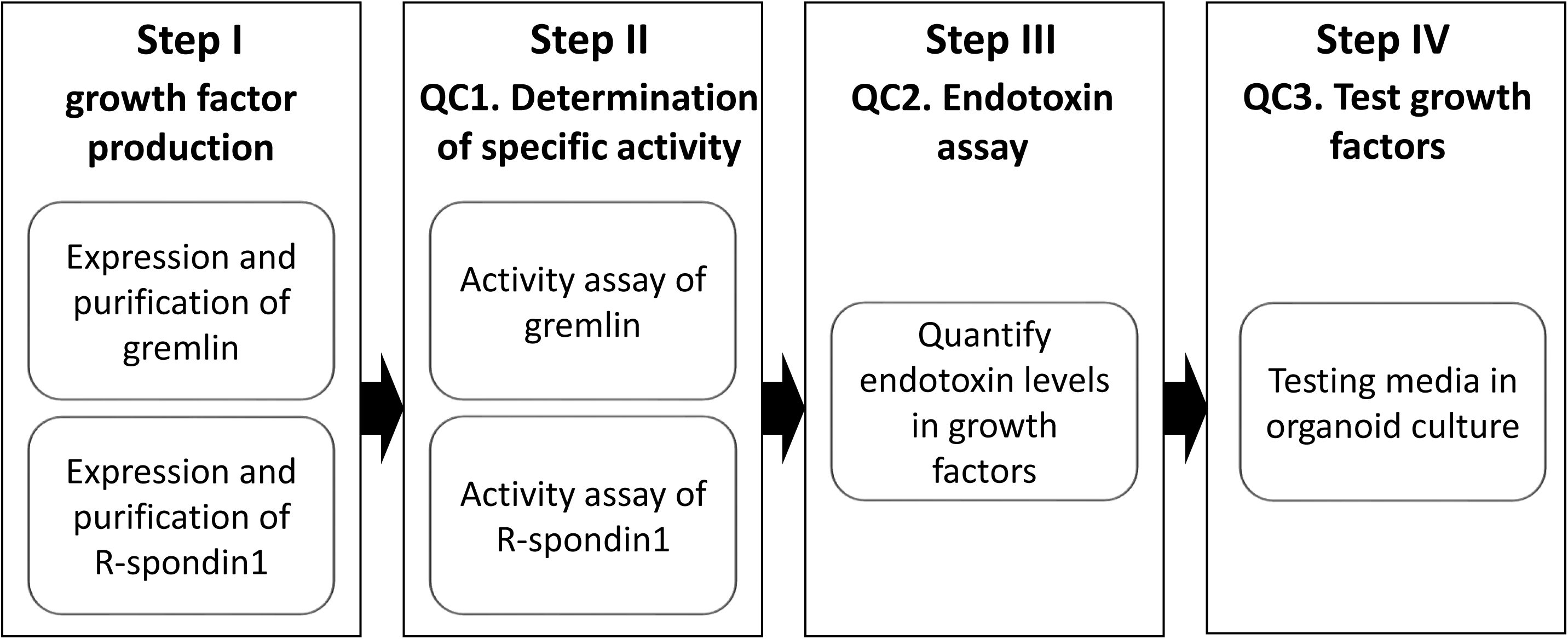
Overview of the procedure. The generation of recombinant R-spondin 1 and Gremlin and subsequent characterisation steps for use in organoid media are as follows: Step I - Recombinant growth factors R-spondin 1 and Gremlin 1 are expressed in bacteria and purified. Step II – Specific activities of Gremlin 1 and R-spondin 1 are determined from cellular reporter assays. Step III – A quality control step is used to ensure minimum endotoxin levels (derived from growth factor expressing bacteria) are present in the preparations using the limulus amoeboid lysate (LAL) assay. Step IV - media formulated with defined activities of R-spondin 1 and Gremlin 1 is tested for the ability to support growth of murine intestinal epithelial organoids. We have further outlined abridged purification protocols for Gremlin 1 and R-spondin 1 in the Materials and Methods, suited to laboratories without access to production-scale chromatography equipment, that produce pure growth factors of sufficient cellular activity for routine organoid culture.

**Figure 2.**
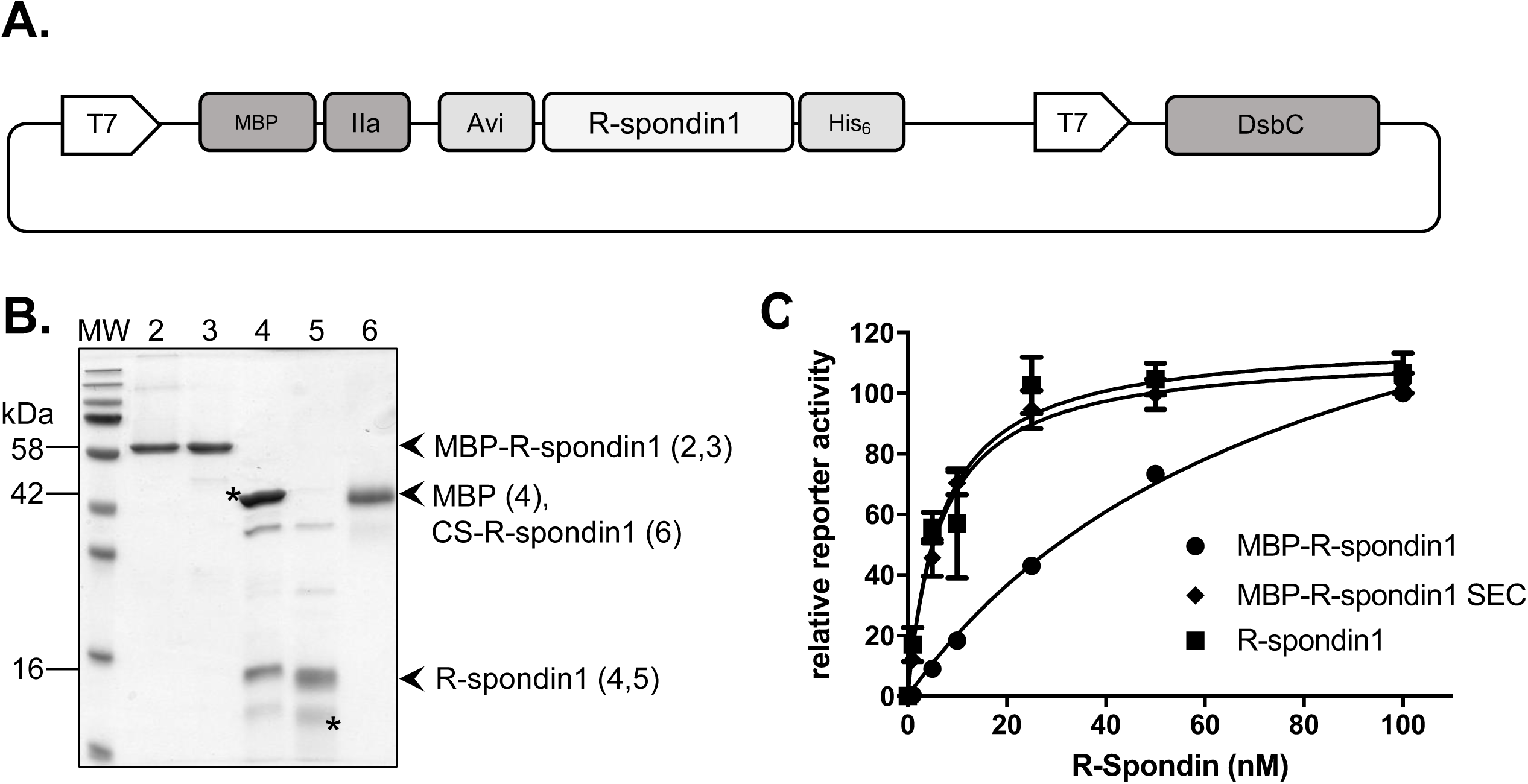
Bacterial expression and production of R-spondin 1. **(A)** Format of the pR-spondin 1 expression vector based on the pETDuet backbone. pR-spondin 1 is a modification of a previously R-spondin 1 expression vector (4). **(B)** SDS-PAGE analysis of fractions from generation of R-spondin 1. For each fraction 2 μg of protein was loaded. Lane numbers (top and in brackets) are: 2 – MBP-R-spondin 1 before SEC, 3 - MBP-R-spondin 1 after SEC, 4 – products of the Thrombin cleavage reaction, 5-final R-spondin 1 preparation and 6 – commercially-sourced R-spondin 1. MW, labelled molecular weight standards. **(C)** Cellular activity of R-spondin 1 fractions measured as the potentiation of Wnt3A-induced SuperTop activity: *circles* - MBP-R-spondin 1 before SEC, WPC50 = 79.3 ± 7.5 nM, *diamonds* - MBP-R-spondin 1 after SEC, WPC50 = 6.6 ± 0.9 nM and *squares* - final R-spondin 1 preparation, WPC50 = 6.8 ± 1.9 nM.

The expression construct, pHAT4-NΔ-Grem1, encoding for a protein with hexa-histidine tag, TEV protease cleavage site and amino acids 72-184 of human Gremlin 1 (Uniprot: O60565), has been described previously ((5); Fig. 3A). Expression of human BMP2 from the bacterial plasmid pBAT-BMP2 and its purification has been detailed elsewhere (5).

**Figure 3.**
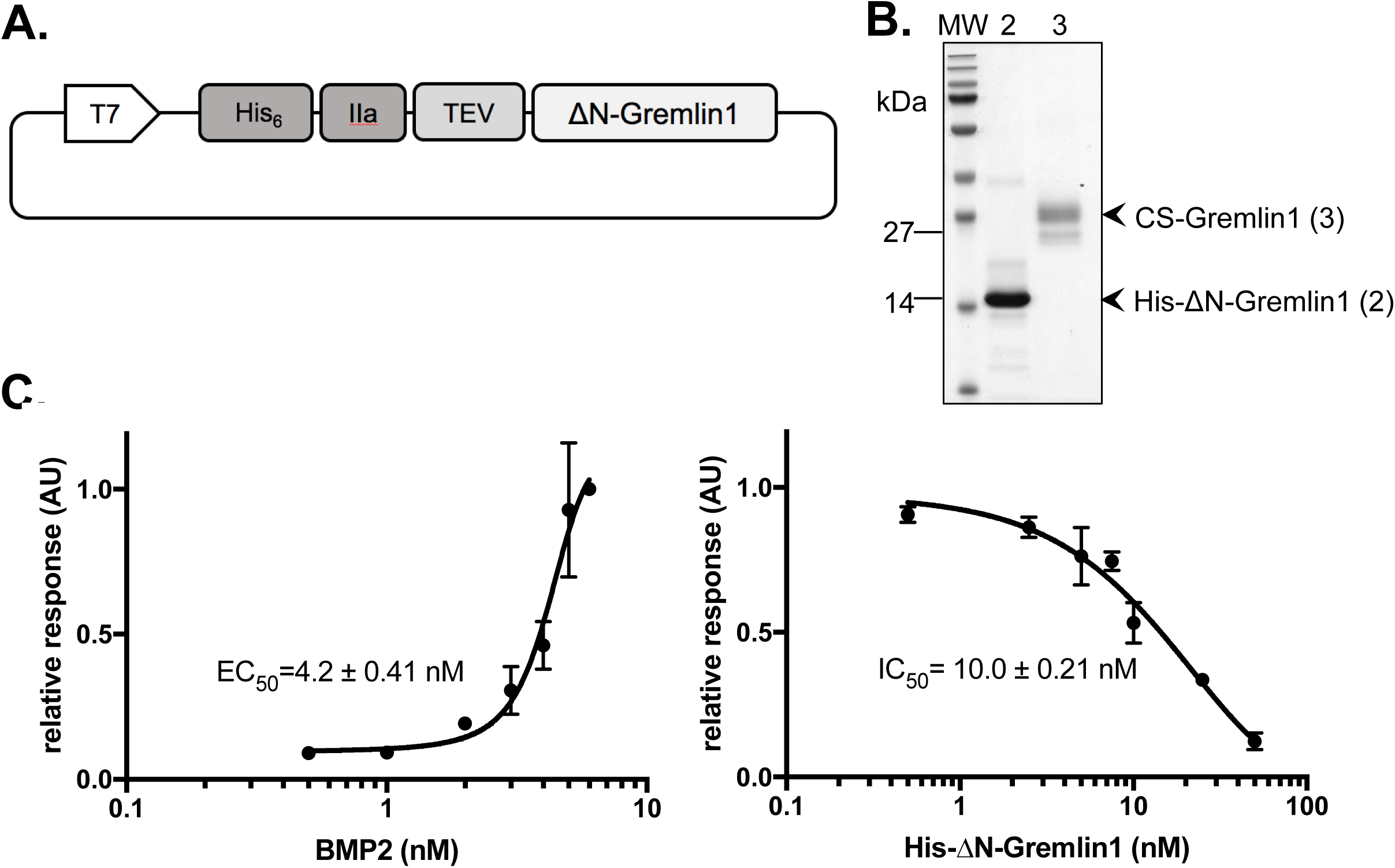
Bacterial expression and production of Gremlin 1. **(A)** Format of the His-ΔN-Gremlin 1 plasmid (5). **(B)** SDS-PAGE analysis of purified His-Δ N-Gremlin 1 (lane 2) and commercially-sourced Gremlin 1 (lane 3), 2 μg of each protein was loaded. MW, labelled molecular weight standards. **(C)** Activity assay for purified Gremlin 1 based on its inhibition of the BMP2-induced alkaline phosphatase in C2C12 cells. *Left graph* – Determination of BMP2 bioactivity to establish BMP2 concentration for use in cellular inhibition assay to evaluate Gremlin 1 bioactivity. *Right graph*: Inhibition of BMP2-induced alkaline phosphatase activity by Gremlin 1. IC_50_ value for Gremlin 1 was determined as 10 ± 0.21 nM.

### Cell lines

HEK293T and C2C12 cells were obtained from ATCC. The cell lines are routinely screened for mycoplasma contamination and validated by short tandem repeat DNA profiling (STR) and DNA fingerprinting as described in the ATCC SDO Workgroup ASN-0002 Standards document (6).

### Organoid media components

Components of the respective media formulations for each organoid type used in the study, listed in Table 1, are as follows: Base media - Advanced DMEM F12 (Gibco) containing 1X penicillin/streptomycin and 10 mM HEPES buffer pH 7.0. Supplements include the following: B27 and N2 supplements (Gibco), N-acetyl-L-cysteine (NAC; Sigma-Aldrich), human gastrin 1 (Sigma-Aldrich), A83-01 (Sigma-Aldrich), SB202190 (Sigma-Aldrich), prostaglandin E2 (PGE2; Sigma-Aldrich), Gastrin I (Sigma-Aldrich), nicotinamide (Sigma-Aldrich) and Primocin (InvivoGen).

**Table 1.** Media components for the culture of organoids. Organoids derived from murine small intestinal epithelial, colon epithelial and mammary gland and human colon epithelia and patient-matched colon carcinoma were formulated as previously described (10, 13) and include optimised concentrations of bacterially-derived R-spondin 1 and Gremlin 1.

| Component | Source of derived organoids |  |  |  |  |
| --- | --- | --- | --- | --- | --- |
|  | murine small intestinal epithelial | murine colon epithelial | murine mammary gland | Human colon epithelia | Human colon carcinoma |
| Base media* | 1X | 1X | 1X | 1X | 1X |
| B27 suppl. | 1X | 1X | 1X | 1X | 1X |
| N2 suppl. | 1X | 1X | 1X | 1X | 1X |
| NAC | 1.25 mM |  | - | 1 mM | 1 mM |
| EGF | 50 µg/ml | 50 µg/ml | - | 50 µg/ml | 50 µg/ml |
| R-spondin 1 | 25 nM | 25 nM | 25 nM | 25 nM | 25 nM |
| Gremlin 1 | 25 nM | 25 nM | 25 nM | 25 nM | 25 nM |
| WCM | - | 0.5X | - | - | 0.5X |
| NRG-1 | - | - | 100 ng.ml | - | - |
| A83-01 | - | - | - | 500 nM | 500 nM |
| SB202190 | - | - | - | 10 µM | 10 µM |
| PGE2 | - | - | - | 10 mM | 10 mM |
| FGF |  |  |  | 100 µg/ml | 100 µg/ml |
| Gastrin I | - | - | - | 10 nM | 10 nM |
| Nicotinamide | - | - | - | 10 mM | 10 mM |
| Primocin | - | - | - | 1X | 1X |

\* Base media is DMEM/F12 media containing 2 mM glutamine, 10 mM HEPES and 0.5 U/ml Penicillin/streptomycin

Commercially-sourced growth factors used in the study are: R-spondin 1 (amino acids 31-263; cat. 4645-RS-025), Gremlin 1 (cat. 5190-GR-050), FGF-2 (cat. 233-FB-010), Neuregulin-1/NRG-1 (cat. 396-HB/CF) and EGF (cat. 236-EG-200), all from R&D Systems. All organoids were grown in growth factor reduced, serum free Matrigel (Fisher Scientific).

### Expression and purification of R-spondin 1

Expression of the R-spondin 1 fusion protein from pR-spondin 1 was carried out in the host *E. coli* strain SHuffle®T7 *E.coli*/K12, following a protocol from the Pioszak laboratory (4). An overnight starter culture was used to inoculate 2 x 1 L of Luria Broth (LB) media in 2 L baffled Erlenmeyer flasks and grown to an OD_600_ of 0.7 with shaking at 200 rpm. Flasks were transferred to a 16°C shaker and expression was induced for 16 hours with 0.4 mM IPTG. Unless otherwise noted, all subsequent steps were carried out at 4°C. Cells were harvested by centrifugation at 4000 x g for 10 minutes and the pellet was resuspended in 10 ml of resuspension buffer (50 mM Tris-HCl pH 7.5 containing 150 mM NaCl and 10% glycerol, protease inhibitor cocktail; Roche) per 1 L culture. Cells were lysed in an Avestin EmulsiFlex C5 homogeniser and lysates were clarified by centrifugation at 48,000 g for 30 min. The supernatant was transferred to a 50 ml conical tube containing 2 ml of a 50% slurry of Ni^2+^-agarose pre-equilibrated with resuspension buffer and absorption of MBP-R-spondin 1 to Ni^2+^-agarose was carried out for 1 h with gentle agitation. Lysate and Ni^2+^-agarose beads were transferred to a 14 cm Econo-Pac column (Bio-Rad), washed thrice with 10 ml of resuspension buffer containing 25 mM imidazole and MBP-R-spondin 1 was eluted in 5 fractions with additions of 1 ml resuspension buffer containing 265 mM imidazole.

Eluted fractions containing MBP-R-spondin 1 were pooled and concentrated using an Amicon Ultra-15 centrifuge filter (Millipore). Buffer exchange into disulphide reshuffling buffer (50 mM Tris base, pH 8.0 containing 150 mM NaCl, 5% glycerol, 1 mM EDTA, 5 mM reduced glutathione and 1 mM oxidized glutathione) was carried out using a PD10 Sephadex G25 desalting column (GE Healthcare). The concentration of the pooled fractions was diluted to 1 mg/ml with disulphide reshuffling buffer and incubated for 12 hours at room temperature. The reshuffled protein was injected onto a HiLoad Superdex 200 pg 16/600 column (GE Healthcare) equilibrated in resuspension buffer and resolved fractions from the size exclusion chromatography (SEC) column containing MBP-R-spondin 1 (identified by SDS-PAGE) were pooled and concentrated using an Amicon Ultra-14 centrifuge filter. Buffer exchange was carried out using a PD10 Sephadex G25 desalting column (GE Healthcare) equilibrated in IEX buffer (50 mM Na^2+^/K^+^ phosphate buffer pH 7.0 containing 5 % glycerol). At this stage, MBP-R-spondin 1 fusion protein can be analysed to determine concentration (using the Bradford protein assay; Bio-Rad) and used in organoid growth media at a concentration of 25 nM (see Results). MBP-R-spondin 1 was incubated with 5 U of Thrombin (Sigma) per mg protein for 8 hours at room temperature and the cleaved R-spondin 1 moiety was resolved using cation exchange chromatography (IEX) with a HiTrap SP HP 5 ml column on an AKTA Purifier equilibrated in IEX buffer (15 mM sodium phosphate buffer containing 5% glycerol) with a linear gradient to 1 M NaCl in IEX buffer. R-spondin 1-containing fractions (identified by SDS-PAGE) were pooled, concentrated and buffer exchanged using a PD10 Sephadex column equilibrated in phosphate buffered saline buffer. The cellular activity of purified R-spondin 1 was determined by Wnt pathway reporter assay (see below).

### Expression and purification of Gremlin 1

Human Gremlin 1 was expressed from a T7-based His-tag vector, His-Δ N-Gremlin 1 (encoding amino acids 72-184 of Gremlin 1), in the *E. coli* strain BL21 (DE3) as follows: cells from glycerol stock were plated on L-agar plates with 100 μg/ml of ampicillin and grown overnight. Cells from the plates were grown in baffled 2 L flasks in 1 L of 2xYT media until an OD_600_ of 0.8 at which point Gremlin1 expression was induced with 400 μg/ml isopropyl β-D-1-thiogalactopyranoside (IPTG) for three hours at 37°C. Cells were harvested by centrifugation for 20 minutes at 4,000 x g and stored frozen at −20C. The frozen pellet from 1 L of Gremlin 1 expressing culture was resuspended in lysis buffer (50 mM Tris base pH 8.0, 2 mM EDTA pH 8.0, 10 mM DTT, 0.5% Triton-X100) and lysed using an Emulsiflex C5 homogeniser. The lysate was incubated for 20 minutes at 25°C with DNase I (Sigma) and 4 mM magnesium chloride and clarified by centrifugation for 30 minutes at 15,000 x g. The supernatant was discarded and the pellet fraction containing insoluble inclusion bodies was washed by resuspension in lysis buffer followed by centrifugation for 30 minutes at 15,000 x g. The pellet was re-washed with lysis buffer containing 1 M NaCl and finally with lysis buffer without Triton-X100. The pellet was resuspended in 5 ml of 100 mM Tris(2-carboxyethyl)phosphine (TCEP), to which 15 ml of 50 mM Tris-HCl pH 8.0, 0.5 mM EDTA containing 8 M guanidine hydrochloride (GndHCl) was added (resulting in a final concentration of 6 M GndHCl and 25 mM TCEP). The resuspended pellet was incubated for 15 minutes at 25°C, clarified by centrifugation (20 min, 15000 x g) and the supernatant diluted to 100 ml with 6 M urea, 20 mM HCl. The solubilised protein solution was then mixed rapidly with 900 ml refolding buffer (50 mM Tris pH 8.0, 1 M 3-[3-(1,1-bisalkyloxyethyl)pyridin-1-yl]propane-1-sulfonate (PPS), 50 mM ethylenediamine, 2 mM EDTA, 2 mM cysteine and 0.2 mM cystine) and left for 7 days at 4°C to allow the correct formation of the disulphide bonds.

After centrifugation (15,000 x g for 30 mins), the pH of the refolded protein solution was adjusted to 7.0 and loaded onto a 5 ml HiTrap SP HP column equilibrated with binding buffer (20 mM Hepes, pH 7.0). Gremlin was eluted using a linear, 75 ml gradient, of 0% to 100% elution buffer (20 mM Hepes, pH 7.0, 1 M NaCl) and peak fractions, analysed by SDS-PAGE, were pooled. At this stage of the purification, Gremlin 1 can be used in small intestinal epithelial organoid media at a concentration of 25 nM.

Further purification of Gremlin 1 proceeds with the addition of acetonitrile and trifluoroacetic acid to a concentration of 10% and 0.1%, respectively. The sample was loaded onto an ACE 5 C8-300 (4.6 × 250 mm) reversed phase column (HiChrom) equilibrated with binding buffer (10% ACN, 0.1% TFA). Proteins were eluted with a linear 20 ml gradient of 20% to 40% ACN containing 0.1% TFA. Peak fractions with pure protein (analysed by SDS-PAGE) are pooled and concentration determined using absorbance at 280 nm with a calculated molar absorption coefficient of 11,460 mol^−1^·cm^−1^. 500 μg Aliquots of 500 μg each were dried under vacuum in a centrifugal concentrator and stored at −80°C. The protein is stable for at least 24 months when stored in this way.

Prior to use, purified Gremlin 1 was reconstituted in PBS buffer at a concentration of ‘1 mg/ml, and confirmed by Cellular activity was determined by alkaline phosphatase assay (see below).

### Cellular reporter assay of R-spondin 1 activity

The cellular activity of MBP-R-spondin and R-spondin 1 fractions were determined by the ability to potentiate Wnt3A-induced pathway activity in HEK293T cells, designated as WPC50 (Wnt pathway activity potentiation constant). Cells were plated onto 24-well tissue culture plates (75,000 cells per well) in DMEM media supplemented with 10% fetal calf serum (Gibco) and transfected with 100 ng SuperTop and 10 ng pRL-SV40P plasmids. Transfected cells were treated with Wnt3A-conditioned media (WCM; produced from Wnt3A-expressing L-cells; ATCC CRL-2646) diluted 1:4 with growth media and varying concentrations of R-spondin 1 (typically 1 – 100 nM). At 16 hours post-treatment, cells were harvested and reporter activities determined with the Promega Stop & Glo kit using a PHERAstar microplate reader. Normalised luciferase activity was determined by dividing firefly luciferase values by the *Renilla* luciferase values. EC50 values for the MBP-R-spondin and R-spondin 1 fractions were calculated using GraphPad Prism Software Version 5. SuperTop activity values for each concentration of R-spondin were the average of 4 individual experimental replicates and are reported with standard deviation (SD).

### Cellular Gremlin 1 activity

The cellular activity of purified Gremlin 1 is defined as the IC_50_ for inhibition of BMP2-induced cellular activity using the alkaline phosphatase assay (ALP)(7). Optimal concentration of BMP2 for use in Gremlin 1 activity assays was determined as follows: C2C12 cells (low passage number is critical) were plated into 96-well plates and cultured in DMEM containing 10 % FBS. To determine optimal concentrations of BMP2 for the Gremlin 1 inhibition assay, a serial dilution of BMP2 (0 nM - 10 nM) was added to the cells and incubated for 48 hrs. Cells were washed with PBS and alkaline phosphatase activity was measured by adding the substrate *p*-nitrophenyl phosphate (pNPP; Sigma Aldrich) and absorbance read at 405 nm using a PHERAstar FS plate reader. The EC_50_ values for BMP2 activity was calculated as 4.2 ± 0.41 nM using Graphpad Prism with a maximal activity at 10.0 nM. Activity values for each concentration of BMP2 were the average of 3 individual experimental replicates reported with SD. The sub-maximal value of 6 nM BMP2 was chosen for use in Gremlin 1 activity assays.

Determination of the Gremlin 1 IC_50_ for inhibition of BMP2-induced cellular activity was carried out as above, using serial dilutions of Gremlin 1 in the presence of 6 nM BMP2. IC_50_were using GraphPad Prism. Activity values for each concentration of BMP2 and were the average of 4 individual experimental replicates reported with SD.

### Endotoxin assay

Levels of endotoxin in the R-spondin 1 and Gremlin 1 preparations was determined using the ToxinSensor system (GenScript) according to manufacturer’s protocol. Values were determined from at least two individual preparations of R-spondin 1 and Gremlin 1 fractions, measured with two technical replicates and reported as average values with SD. We have set a maximum endotoxin threshold value of 0.5 EU/ml in organoid media, below the Lowest Observed Cellular Effect (LOCE) threshold for detectable cellular cytokine production and NF-kB activation, previously determined in cell culture assays (8). Each value

### Derivation of organoid cultures

Murine small intestinal epithelial organoids and colon epithelial organoids (murine and human) were derived according to Sato et al. 2013 (9) and Sato et al. 2011 (10), respectively. Murine mammary gland organoids were derived according to Nguyen-Ngoc et al. 2011 (11). A summary of all media formulations used in the study is shown in Table 1. All derived organoid cultures were grown in 48-well CellStar plates (Sigma-Aldrich), embedded in a 25 μL droplet of Matrigel (Corning), cultured in 200 μL of the corresponding organoid media in an incubator at 37°C and 5% CO_2_.

### Optimisation of R-spondin 1 and Gremlin 1 concentration in media formulations

Optimisation of R-spondin 1 and Gremlin 1 concentrations for use in media for culture of murine small intestinal epithelial organoids were determined by growing organoids in individual wells with variable concentrations of R-spondin 1 (0 - 100 nM) or Gremlin 1 (0 – 25 nM) in otherwise fully-supplemented organoid growth media. Growth characteristics (multiplicity and crypt number) of organoids were tested 8 days post-plating. Organoid multiplicity was determined by determining the average count of organoids in each condition form 8 different wells, reported with SD. Organoid crypt number was determined as the average number of crypts from 10 organoids in individual wells. Crypt numbers were subsequently averaged across 8 wells, reported with SD.

## Results

The R-spondin family and the Noggin and Gremlin BMP inhibitor families are essential growth factors for culturing organoids derived from a number of epithelial tissues. We report here a key refinement in the culture of organoids – the use of highly pure preparations of R-spondin 1 and Gremlin 1 of defined cellular activities derived from bacterial expression

### Production of R-spondin 1

The expression vector for R-spondin 1 (Fig. 2A) has been modified from a previous report (12) to include the coding sequence for the 9 amino acid Avi-tag fused to the N-terminus of the R-spondin 1 that confers enhanced solubility and specific activity of the expressed protein.

R-spondin activity is particularly sensitive to the correct configuration of disulphide linkages. In addition to the R-spondin 1 expression cassette, pR-spondin 1 drives expression of *DsbC,* a disulphide isomerase that promotes the correct configuration of disulphide linkages. We express R-spondin 1 in the NEB Shuffle T7 *E. coli* expression strain that also expresses additional *DsbC* in the cytoplasm. Moreover, following lysis of expressing bacteria and batch purification by Nickel-NTA agarose we subject the expressed R-spondin 1 to an *in vitro* disulphide shuffling step in the presence of reduced and oxidised glutathione. Nonetheless, a subsequent SEC step removes approximately 50% of the inactive, insoluble R-spondin 1 protein from preparation yielding pure MBP-R-spondin 1 that migrates as a single band of protein by SDS-PAGE at the expected molecular mass of 58 kDa (Fig. 2B). Cellular activities of the MBP-R-spondin 1 fractions (WPC50s) increase greater than 10-fold with SEC, from 79.3 nM to 6.6 nM (Fig. 2B). The purified MBP-R-spondin can be used to supplement organoid media for routine use at a concentration of 25 nM.

Removal of the MBP moiety by Thrombin cleavage and cation exchange chromatography produces pure R-spondin 1 with a cellular activity of 6.8 nM, similar to MBP-R-spondin 1 after SEC. R-spondin 1 migrates on SDS-PAGE gels as a single band at the expected molecular mass of 16 kDa (Fig. 2B). By contrast, commercially-sourced R-spondin 1 (CS-R-spondin 1) migrates as a 40 kDa protein instead of the expected 26 kDa (predicted from the sequence), presumably due to glycosylation. Note that the CS-R-spondin retains 118 amino acids at the C-terminus not present in bacterially-derived R-spondin 1.

A typical yield of purified MBP-R-spondin 1 and R-spondin 1 is approximately 2.5 mg and 1 mg of protein per L of Luria broth; for culture of murine small intestinal epithelial organoids, 2.5 mg or 1 mg of the respective proteins is sufficient to supplement approximately 3 L of organoid media (at a concentration of 25 nM).

Typical levels of contaminating endotoxin in the final MBP-R-spondin 1 and R-spondin 1 preparations are similar to CS-R-spondin 1 and > 20-fold less than the LOCE threshold value of 0.5 EU/ml endotoxin when diluted into organoid culture media (Table 2).

**Table 2.**
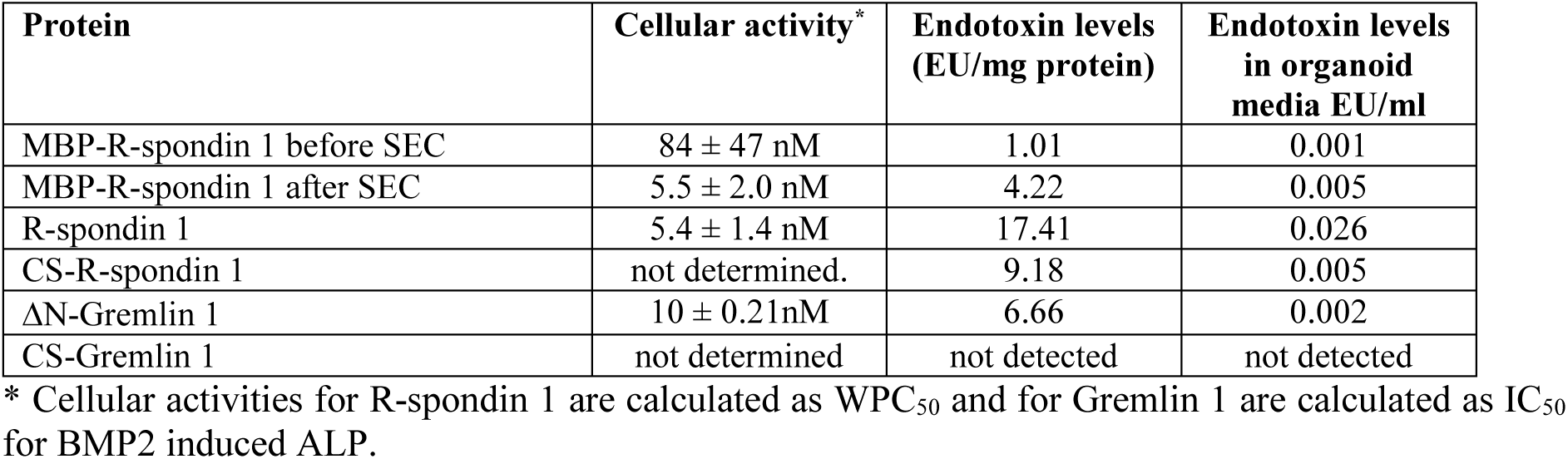
Cellular activities and endotoxin levels in R-spondin 1 and Gremlin 1 fractions.

### Production of Gremlin 1

The vector for bacterial expression of recombinant human Gremlin 1 (His-Δ N-Gremlin 1) and the purification procedure has previously been described (5). Bacterially-expressed recombinant Gremlin 1 associates with inclusion bodies requiring suspension of the protein under denaturing conditions and a refolding/disulphide shuffling step for generation of natively-folded protein. Subsequent anion exchange chromatography leads to an essentially pure preparation of Gremlin 1 that can be used in organoid media formulations at a concentration of 25 nM.

A final reverse phase chromatography step resolves a highly pure fraction of Gremlin 1 shown by a single band on SDS-PAGE migrating at the expected molecular mass of 14 kDa (Fig. 3B). By contrast, the 30 kDa commercially-sourced Gremlin (CS-Gremlin) protein, consisting of amino acids 25-284, migrates as two bands on SDS-PAGE gels, one at approximately 27 kDa and the other at the expected 30 kDa (Fig.3B). The reduced mobility and diffuseness of CS-Gremlin migration is typical of glycosylated proteins derived from eukaryotic expression systems.

The cellular activity of bacterially-produce Gremlin-1 is measured as the IC_50_ for the inhibition of BMP2-induced alkaline phosphatase expression in C2C12 cells per μM protein – typical values for bacterially-expressed Gremlin 1 are (Fig. 3B; Table 2). Endotoxin levels are approximately 250-fold lower that the LOCE threshold (Table 2). A typical yield of pure Gremlin 1 is approximately 10 mg per L of 2-YT broth culture. For the culture of small intestinal epithelial organoids, 10 mg of Gremlin 1 is sufficient for approximately 28 L of media at a concentration of 25 nM.

### Use of R-spondin 1 and Gremlin 1in organoid media formulation

We compared the ability of bacterially-derived R-spondin 1 and Gremlin 1 with the corresponding commercially-sourced growth factors, CS-R-spondin 1 and CS-Gremlin 1 to sustain growth of murine small intestinal epithelial organoids. We find that the activities of R-spondin 1 and CS-R-spondin 1 are similar, each sufficient at a minimum concentration of 5 nM to fully support the growth of organoid cultures (Fig. 4A). We analysed crypt multiplicity in organoids as a more sensitive probe of R-spondin 1 activity. The addition of bacterially-derived R-spondin 1 or CS-R-spondin 1 to organoid media display similar activities after 8 days of growth with a minimum concentration of 25 nM yielding the maximal number of 15 crypts per organoid.

**Figure 4.**
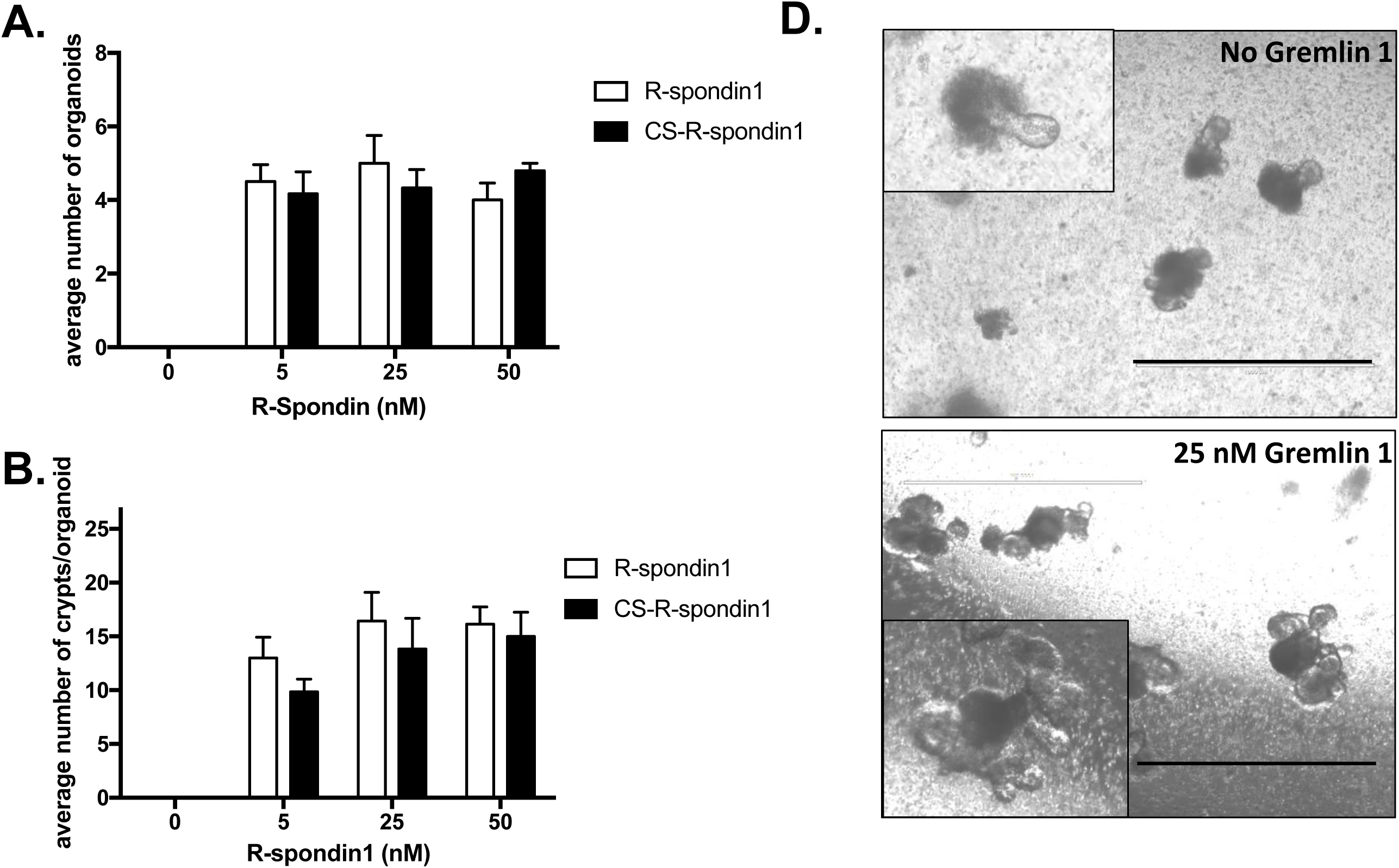
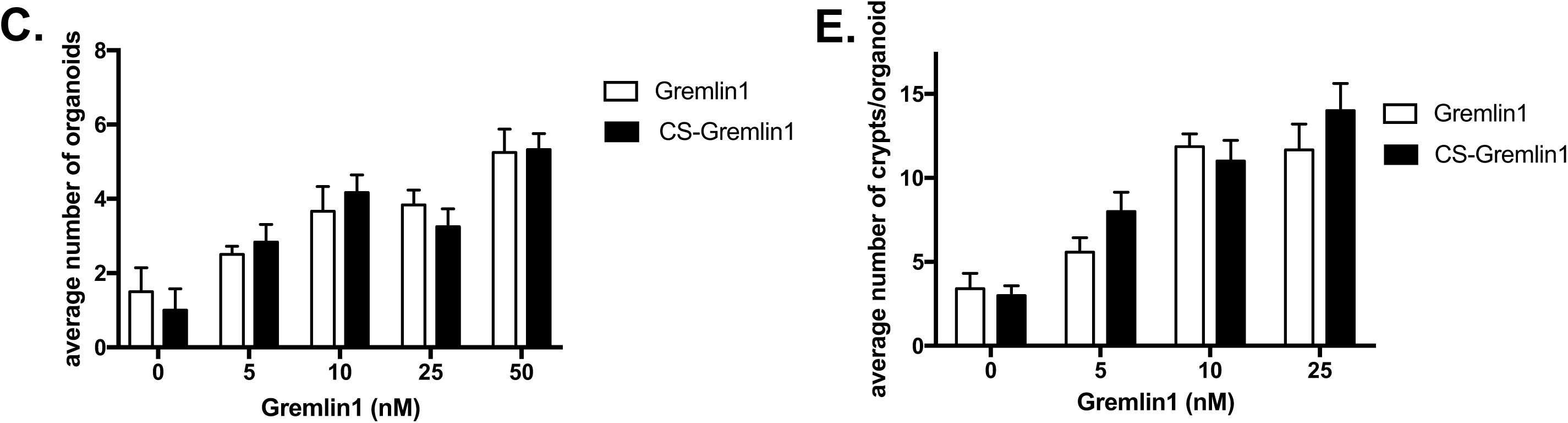
Optimal concentrations of recombinant R-spondin 1 and Gremlin 1 for organoid growth. **(A)** Media formulated with R-spondin 1 or CS-R-spondin 1 equally sustains growth of murine small intestinal epithelial organoids at a minimal concentration of 5 nM. **(B)** Organoid growth media supplemented with either 5 nM R-spondin 1 or 25 nM CS-R-spondin 1 achieves a maximum of crypts per organoid after 8 days growth. **(C)** Gremlin 1 or CS-Gremlin 1 supplemented media equally sustains maximal organoid growth at a minimum concentration of 10 nM. Note that Gremlin 1 is not strictly required in growth media as some organoid growth is maintained in its absence. (D) Addition of 10 nM of either Gremlin 1 or CS-Gremlin 1 is sufficient for a maximum of crypts per organoid after 8 days growth.

The addition of Gremlin to organoid media is not strictly required for growth; we find that media formulations that lack Gremlin 1 and other BMP signalling inhibitors (such as Noggin) can sustain growth of murine intestinal epithelial organoids for up to 2-3 passages as previously described (13). Gremlin 1 or CS-Gremlin 1 have similar activities in organoid media and at 10 nM either protein sustains long-term organoid cultures (Fig. 4C). Analysis of crypt multiplicity after 8 days growth indicates that a minimum of 25 nM Gremlin 1 or CS-Gremlin 1 yields the maximum number of crypts per organoid.

### Application to organoids derived from a range of tissue types

We tested the ability of bacterially-derived R-spondin 1 and NΔ-Gremlin 1 in corresponding media formulations (Table 1) to support sustained culture of organoids derived from a number of murine tissues including colon epithelia, APC^min/-^ small intestinal tumours and mammary gland (Figs 5A, B and C) as well as organoids derived from clinical biopsies of human colon epithelia and patient-matched colon cancer tumours (Fig. 5D; media formulations in Table 1). In all cases, media containing bacterially-expressed R-spondin 1 and NΔ-Gremlin 1 supported robust organoid growth indicating wide application of our protocols.

**Figure 5.**
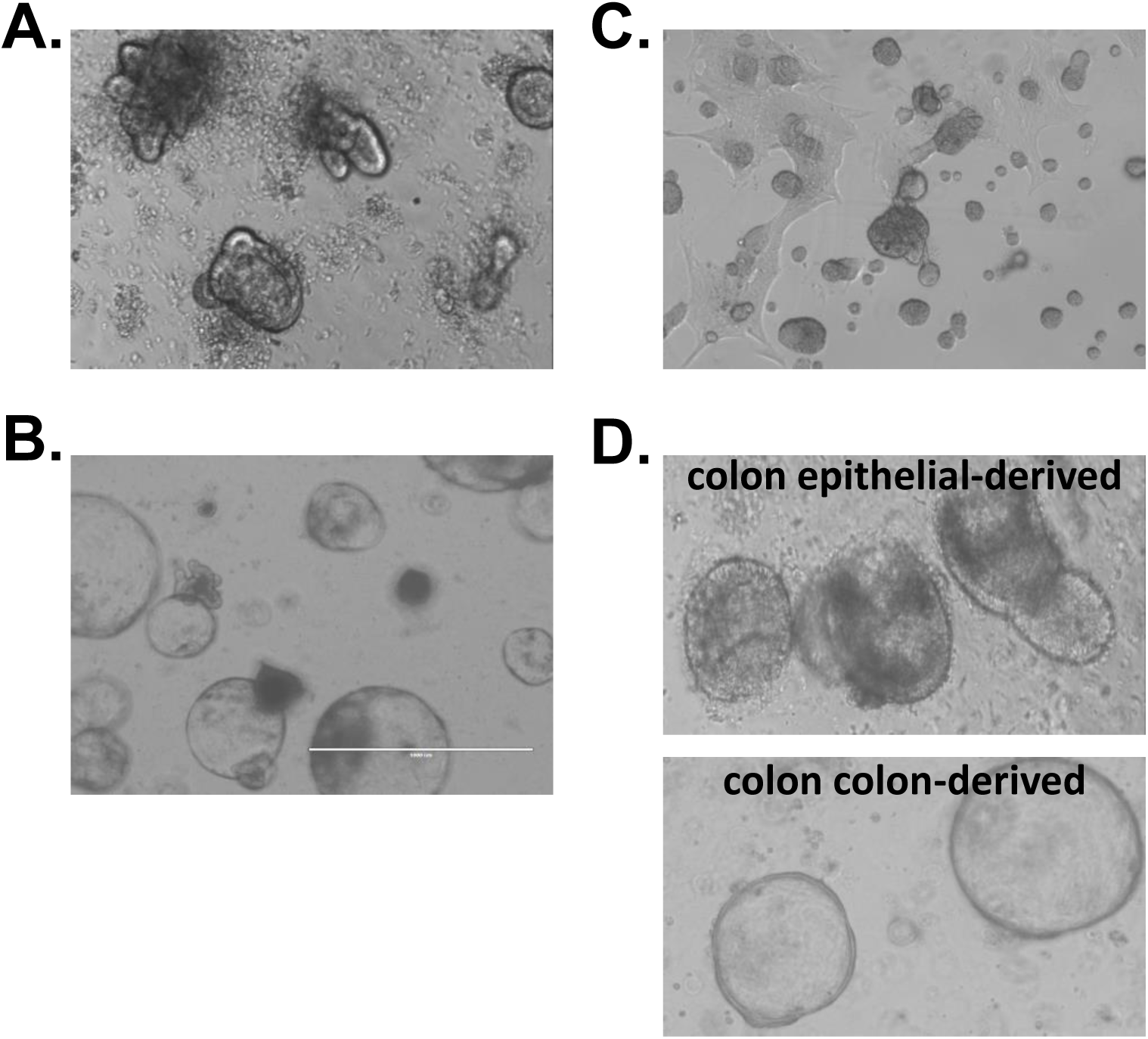
Growth media formulated with R-spondin 1 or Gremlin 1 can sustain growth of a organoid types. The corresponding media formulations (Table 1) can be used to culture organoids derived from: **(A)** murine colon epithelia, **(B)** murine APC^min/-^ small intestinal tumours, **(C)** murine mammary gland and (**D**) human biopsy-derived colon epithelia (top panel) and human colon adenocarcinoma (bottom panel).

## Discussion

Organoid culture, though still being refined for some epithelial tissues, is fast becoming an essential laboratory tool. Current implementation of organoid technology has graduated from proof-in-principle studies to hypothesis testing in the field of epithelial autonomous physiology, research into the molecular underpinnings of genetic diseases such as cancer (2) and cystic fibrosis (14) and the recapitulation of epithelial and tumour microenvironments (15) and host-virus interactions (16). We report here the production of highly pure Gremlin 1 and R-spondin 1, of known cellular activities, derived from bacterial expression for use in organoid media. The protocol we outline has three major advantages over conventional use of Gremlin 1 and R-spondin 1 from conditioned media or commercial sources: (i) highly pure R-spondin 1 and Gremlin 1 is devoid of impurities such as serum-derived growth factors; (ii) the use of precise cellular activities of the growth factors minimising the effects of batch-to-batch variability. ii) a substantial reduction in cost of organoid media. The protocol we outline for the generation if media containing recombinant R-spondin 1 and Gremlin 1 is therefore a leap forward in the ability to generate reproducible growth conditions for both routine organoid culture and large-scale applications such as genetic and chemical screens or clinical tissue biobanking previously reported for limited sample sets of colon cancers (2, 3). We have further demonstrated the applicability of bacterially-derived growth factor production for the culture of organoids derived from a range of tissues including murine small intestine and colon epithelia, mammary gland and human colon epithelia and colon cancer biopsies. We envision that the use of organoid media formulations with defined cellular activities of growth factors will spur the emergence of new applications for organoids that, in particular, require large-scale, cost-effective culture.

One caveat in the implementation of our protocols is the requirement for an animal cell-derived surrogate extracellular matrix. Currently, supports such as Matrigel, are produced from Engelbreth-Holm-Swarm mouse sarcoma cells and may contain serum impurities that impact organoid growth. However, synthetic matrices supporting have recently been designed and show promise for supporting growth (17). Further development of synthetic matrices and their use in conjunction with our protocols will ultimately lead to a fully-defined organoid culture system.

## Author contributions

MU designed and carried out experiments, interpreted data and generated figures. KR generated recombinant Gremlin 1, HR and TF contributed organoids and reagents to the study. MH supervised the study and designed experiments. MAD supervised the study, designed experiments, interpreted data and drafted the manuscript.

## Data Availability

The data generated during and/or analysed during the current study are available from the corresponding authors.

